# A novel clonal and axenic isolation pipeline for the human gut protist *Blastocystis* reveals within-subtype strain coexistence

**DOI:** 10.64898/2026.10.19.760120

**Authors:** Fredrick Leon, Johnathan B. Shih, Lydia E. Keller, Shobini Palaniappan, Lucie Degaetano, Gabriela A. Nine, Abigail L. Lind

**Affiliations:** School of Biological Sciences, Georgia Institute of Technology, Atlanta, Georgia, USA; Interdisciplinary Graduate Program in Quantitative Bioscience, Georgia Institute of Technology, Atlanta, Georgia, USA

## Abstract

*Blastocystis* is the most prevalent protist in the human gut microbiome, but its study is impeded by limitations in existing cultivation methods that hinder routine axenic and clonal isolation. These barriers have left open the question of whether multiple strains of a protist species can coexist within a single host, as observed within the bacterial gut microbiota. To address this, we introduce egg-based monophasic (EBM) medium, a novel optically clear liquid medium that supports exponential growth of *Blastocystis* from low inoculum densities in standard tissue-culture plates and flasks, a capability not supported by existing media formulations tested here. Using EBM medium, we established a modular axenization and clonal isolation pipeline and recovered axenic and clonally isolated cultures representing *Blastocystis* subtypes 1–4 from human and non-human primate hosts. To investigate within-host strain diversity, we sequenced a subtype 1 culture before and after clonal isolation. The non-clonal culture yielded a 21 Mb assembly with gene copy distribution and allelic polymorphism patterns consistent with multiple strains, whereas its clonal derivative yielded a 15 Mb assembly with patterns consistent with a single strain. These results indicate that the original culture contained multiple closely related strains co-colonizing one host. We identified the same genomic signature in published xenic cultures established from single human donors, supporting the presence of strain-level diversity within *Blastocystis* subtypes. Our pipeline provides a scalable platform for *Blastocystis* cultivation, imaging, and clonal isolation, enabling investigation of strain-level diversity that may be obscured in non-clonal cultures.

**Importance:** *Blastocystis* is the most prevalent protist in the human gut, yet experimental investigation remains limited by a difficulty in obtaining single-microbe and genetically defined cultures. Existing methods make it difficult to separate the protist from associated bacteria or recover individual lineages from mixed cultures. We developed a liquid growth medium compatible with standard tissue culture plates, live-cell imaging, and a modular workflow for producing axenic and clonal isolates. We use this pipeline to create axenic and clonal isolates across four common subtypes. Applying this workflow exposed a mixture of closely related strains in a single-subtype culture, a finding supported by analyses of independent published cultures. This culturing pipeline enables controlled investigation of strain-level diversity, and provides a new experimental framework for the study of *Blastocystis* biology and its interactions with the gut microbiome.

## Introduction

The vertebrate gut microbiome is an ecologically diverse mixture of microbes from all domains of life, including bacteria, archaea, viruses, and microbial eukaryotes like protists and fungi^1^. These microbes interact with each other and their host, shaping host phenotypes and contributing to host health and disease^1–3^. Substantial progress has been made in characterizing gut bacteria at the species and strain levels, revealing differences in gene content and host-relevant phenotypes, even among strains of the same species. For example, strains of *Eggerthella lenta* vary in the presence of the cardiac glycoside reductase operon, which determines whether a host’s microbiome inactivates the drug digoxin^4,5^. Further, *Mediterraneibacter gnavus* comprises distinct clades that differ in their association with inflammatory bowel disease^6^. Strain-level variation is also structured within individual hosts, as multiple strains of a single bacterial species can coexist at dynamically stable abundances within the same individual^7^.

Considerably less is known about strain-level diversity among gut eukaryotes. *Blastocystis*, the most prevalent human gut protist^8,9^, is a genus of obligately anaerobic stramenopiles divided into 18S rRNA-defined subtypes (STs), with genetic divergence comparable to interspecies differences in other protists^10,11^. It occurs in up to 50% of individuals in industrialized populations and up to 95% in non-industrialized populations^8,9^, yet its role in the gut remains unresolved. Historically regarded as a possible pathogen^12^, it has more recently been associated with markers of health, including lower gut inflammatory markers and favorable cardiometabolic outcomes^9,13–15^. These associations are correlational and vary across cohorts and STs, and their direction of causality remains unclear. Whether *Blastocystis* phenotypes depend on ST, variation among strains within a ST, or interactions with the surrounding microbial community remains difficult to determine without experimental systems that enable controlled comparisons among individual lineages.

Progress in developing such systems has been limited by cultivation challenges. *Blastocystis* nutritional requirements remain incompletely defined, and no chemically defined medium is available^16^. Experimental studies depend on relatively few laboratory isolates^15,17,18^. Axenic cultures, free of detectable contaminating microbes, permit investigation of protist-specific phenotypes; clonal cultures, derived from a single founder, permit comparisons in a defined genetic background and enable genetic manipulation. Xenic cultures can be established from feces, but published axenization workflows are labor-intensive, may take months, and often fail^16,19,20^. Clonal isolation is even less routine and has relied on agar-based methods or micromanipulation^16,21–23^.

Another bottleneck is the limited capacity of existing media to support exponential growth from low inocula. Serum-supplemented mammalian cell-culture media have supported *Blastocystis* cultivation, but robust exponential growth at the low cell densities needed for isolation has not been established^24^. To our knowledge, all published exponential growth curves for *Blastocystis* used biphasic egg medium^25–28^, whose solid egg slant complicates use in microplates and live-cell imaging. A liquid medium compatible with low-density growth could therefore improve both cultivation and isolation.

These technical challenges leave a basic ecological question unanswered: can distinct strains of a single *Blastocystis* ST coexist within one host? Although co-occurrence of multiple STs is documented^29^, their high genetic divergence does not establish coexistence among strains. Within-species strain coexistence is common and stable in bacteria^7^, but whether similar patterns occur among gut eukaryotes, or whether closely related lineages tend to exclude one another remains unclear. Addressing this question requires methods that separate individual strains from mixed cultures.

Here, we describe egg-based monophasic (EBM) medium, a clear liquid formulation that supports exponential *Blastocystis* growth at low inoculum densities and enables visual screening of microplate cultures. We use EBM medium in a modular pipeline for axenization and clonal isolation, recovering isolates from ST1– ST4 and benchmarking growth against existing media. We then apply clonal isolation to an ST1 culture from a single host and uncover genomic evidence for coexisting strains that conventional subtype assignment missed. Finally, we examine published sequence data for similar strain-mixture signatures. This work couples a broadly applicable cultivation method to a biological demonstration of the diversity hidden within subtype-defined cultures.

## Results

### A Liquid Cultivation Platform Supports Axenization and Clonal Isolation Across Blastocystis *subtypes*

The American Type Culture Collection (ATCC) recommends maintaining *Blastocystis* in biphasic egg medium comprising a solid egg slant overlaid by Stone’s Modification of Locke’s Solution (hereafter, Locke’s) supplemented with heat-inactivated horse serum (HIHS; ATCC Medium 1671; Fig. 1A). Although this formulation supports *Blastocystis* growth, its solid phase complicates microplate screening and imaging^21,22^. Moreover, few *Blastocystis* strains are axenic^16,19,30^ and verified as clonal^21,22^. The ATCC contains eight axenic *Blastocystis* cultures isolated over 40 years ago. Only three are available (Nand II-ST1, JDR-ST3, and BT1-ST4; Table S1), and none are described as clonal. These limitations constrain the availability of well-characterized isolates for experimental studies.

**Figure 1:**
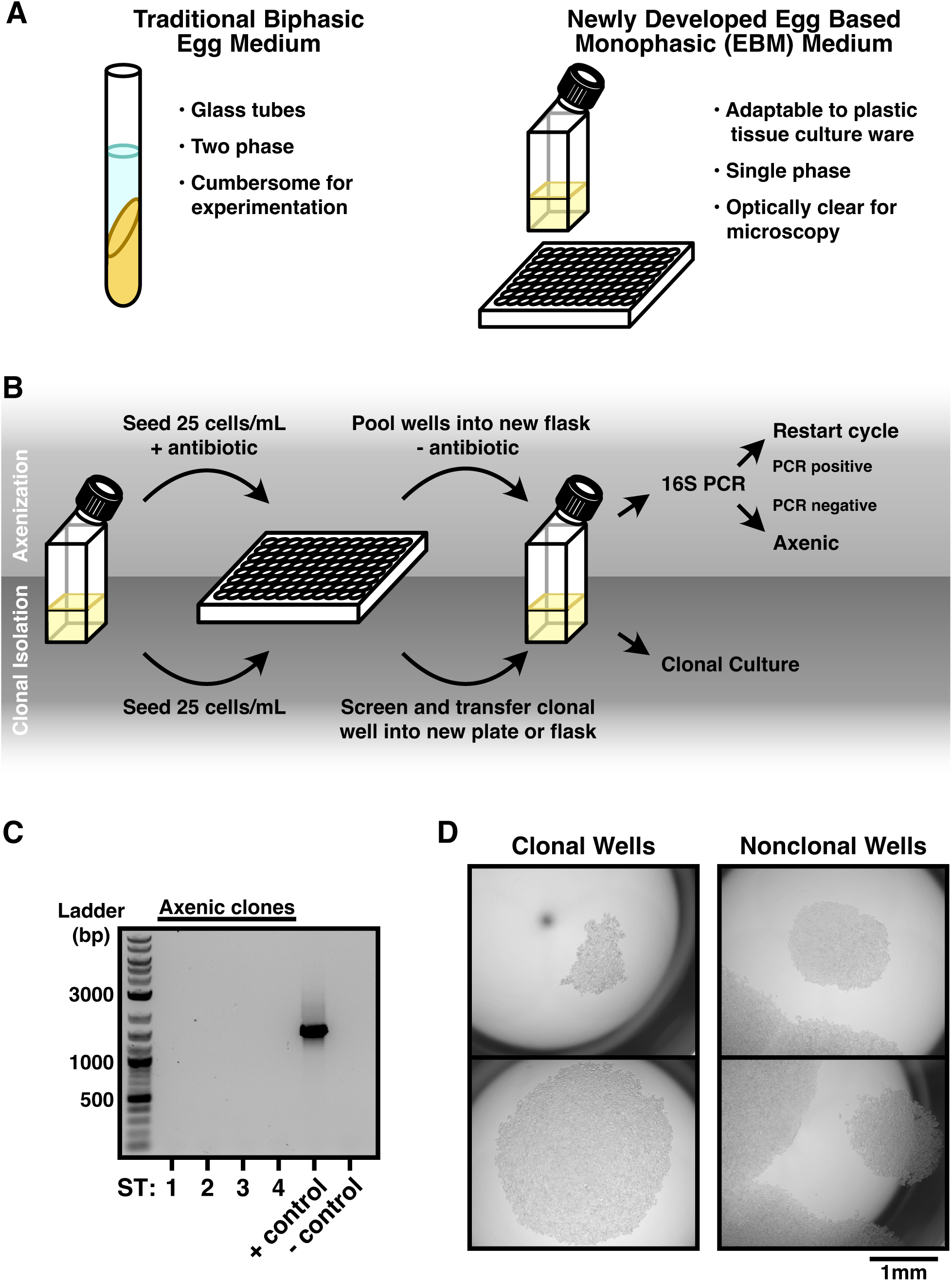
Axenization and clonal isolation of *Blastocystis* primary isolates. **(A)** Comparison of cultivation methods for *Blastocystis*. Biphasic egg media (ATCC Medium 1671) consists of glass tubes with solid egg slants with a liquid overlay. Egg-based monophasic media (EBM) is a monophasic liquid medium comprising egg yolk in a buffered saline solution supplemented with heat-inactivated horse serum. EBM can be utilized in plastic culture plates. **(B)** Axenization and clonal isolation pipelines. **(C)** Gel electrophoresis of 16S PCR products from axenic and clonal *Blastocystis* STs 1-4. The positive control is extracted DNA from a pure *M. gnavus* culture, and the negative control is sterile media. **(D)** Representative images of colonies of *Blastocystis* cells after serial dilution into 96 well plates and growth for 5-7 days. Nonclonal wells have multiple visible colonies.

To address these limitations, we developed a liquid medium, termed Egg Based Monophasic (EBM) medium, that contains components similar to those in traditional biphasic egg medium but is adaptable to standard tissue culture materials. EBM medium contains egg yolk solubilized in Locke’s with Tween 80, supplemented with HIHS and reduced with sodium thioglycolate (Fig. 1A). It is compatible with standard tissue-culture plates and flasks and supports screening in 96-well plates. Tween 80 renders the medium optically clear, enabling live-cell microscopy, including time-lapse imaging of successive *Blastocystis* cell divisions (Movie S1).

We evaluated several media components to optimize growth. The addition of 0.5 mg/mL sodium thioglycolate increased cell density 1.3-fold relative to medium without a reducing agent (*P* = 0.0007, two-way ANOVA; Fig. S1A, Table S2). In contrast, L-cysteine, another commonly used reducing agent, negatively affected growth of the ST1 isolate tested, as did brain heart infusion and tryptic soy broth, which are commonly used in cultivating gut bacteria (Fig. S1A,B, Table S2).

Using EBM, we established two independently deployable modules: axenization by dilution under antibiotic selection and clonal isolation by dilution without antibiotics (Fig. 1B). Both used 96-well plates seeded at 25 cells/mL (2.5 cells per 100-µL well); this density was chosen because plating efficiency decreased at lower inocula (Fig. S1C; Table S2) and after antibiotic exposure. For axenization, candidates were expanded and tested for bacterial contamination by 16S rRNA gene PCR and growth on nutrient-rich agar (Fig. 1B,C); cultures negative by both assays were classified as axenic. For clonal isolation, individual wells were screened for a single visible *Blastocystis* growth focus (Fig. 1B,D), and wells with multiple foci were excluded. Full-length 18S rRNA gene sequencing provided an additional screen for within-culture sequence heterogeneity (see Methods). The order of the two modules can be reversed to suit the starting culture.

With this pipeline, we obtained axenic and clonally isolated cultures representing ST1–ST4 from human and nonhuman-primate fecal samples (Fig. 1C; Table 1; Table S3): ST1 from a Wolf’s guenon, ST2 from a human donor, ST3 from an orangutan, and ST4 from a ring-tailed lemur.

**Table 1.** Strain information for axenic and clonal strains isolated in this study.

| Assigned Subtype | Host species | Isolation Site |
| --- | --- | --- |
| ST1 | *Cercopithecus wolfi* (Wolf's guenon) | Zoo Atlanta |
| ST2 | *Homo sapiens* (Human) | Human Donor |
| ST3 | *Pongo abelii* (Orangutan) | Zoo Atlanta |
| ST4 | *Lemur catta* (Ringtail Lemur) | Zoo Atlanta |

### EBM Medium Supports Robust Growth of Isolates Spanning Four Subtypes

We compared the growth of an axenic and clonal ST1 isolate in EBM medium with and without Tween 80, biphasic egg medium, and Iscove’s Modified Dulbecco’s Medium (IMDM) supplemented with 10% HIHS (Fig. 2A). IMDM has previously been used for *Blastocystis* cultivation and for agar-based isolation protocols^15,21,22,24,31,32^. However, reported doubling times in IMDM range from 21 to 83 hours, require high seeding densities, and published studies have not established exponential growth from low seeding densities^24,33^. To our knowledge, biphasic egg medium is the only published medium shown to support exponential growth of multiple STs^16,19,24–28^, though it cannot be used for clonal isolation procedures due to its solid egg slant.

**Figure 2:**
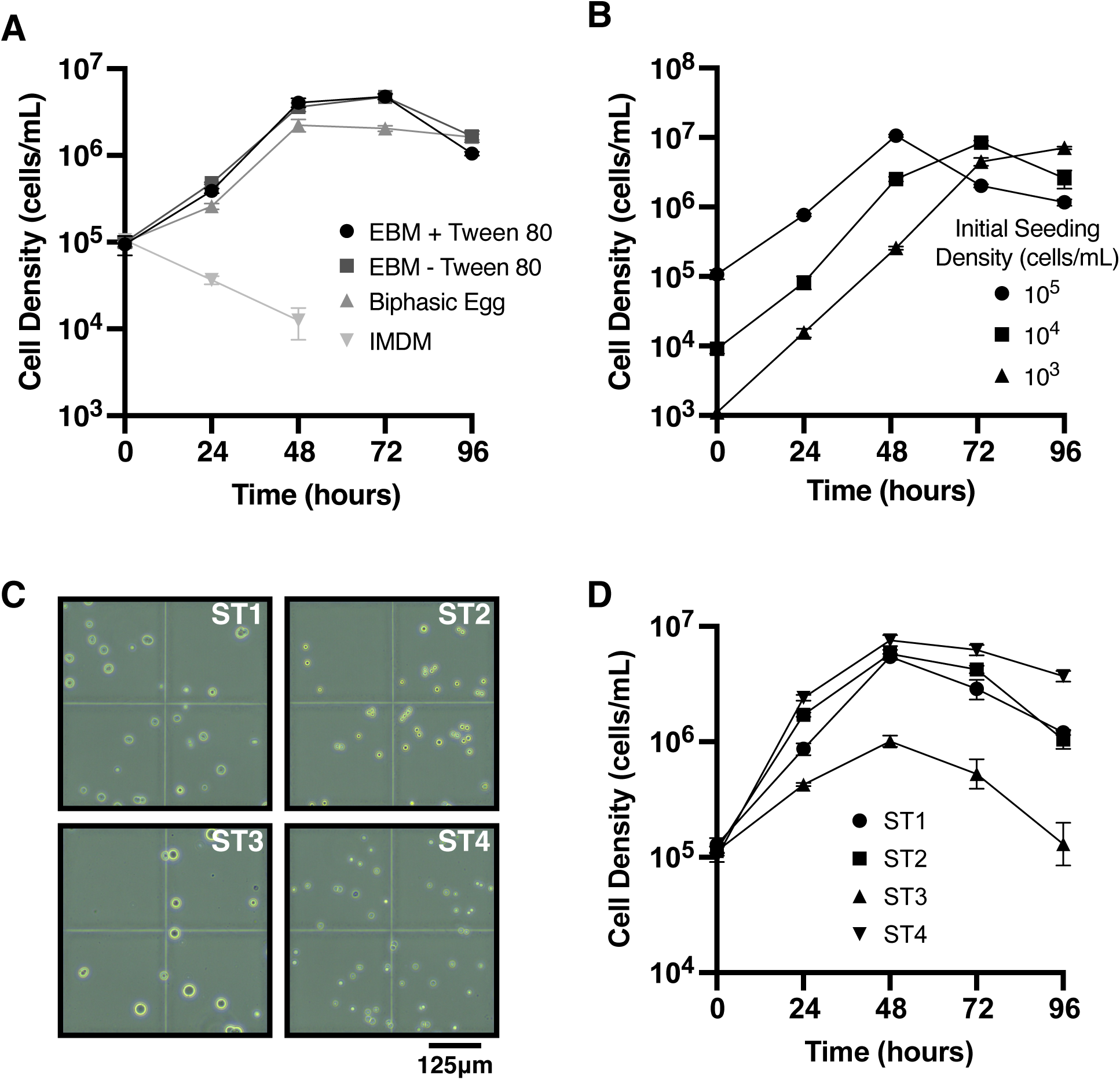
Egg based monophasic medium supports the robust growth of multiple STs. **(A)** *Blastocystis* ST1 cell density across a 96 hour period in EBM medium with and without Tween 80, in biphasic egg medium, and in IMDM. IMDM data stops at 48 hours as zero values cannot be displayed in log scale. Growth curves in EBM medium and IMDM were conducted in flasks to best reflect culture maintenance conditions. **(B)** *Blastocystis* ST1 cell density measured from cultures seeded at variable seeding densities at 10^5^, 10^4^, and 10^3^ seeding densities. Exponential growth is achieved for each initial seeding density tested. Growth curves were conducted in 24 well plates with 1 mL of inoculated EBM medium. **(C)** Axenic and clonal *Blastocystis* cultures, from STs 1-4. Images demonstrate optical clarity of the EMB medium (with Tween 80) and morphological differences between isolated STs. Images were taken via haemocytometer from the 24 hour time point of (D) growth curves. **(D)** *Blastocystis* STs 1-4 grown at 96 hours, with cell densities measured every 24 hours in EBM medium in 24 well plates.

All media were pre-reduced for 48 h and supplemented with sodium thioglycolate except IMDM, which was tested in its previously used formulation^24^. We measured cell density over 96 hours, starting at 10^5^ cells/mL. At 24 hours, densities in both EBM formulations were at least 1.5-fold higher than in biphasic medium (*P* ≤ 0.012; two-way ANOVA; Fig. 2A; Table S2). Their modeled carrying capacities were approximately 2.3-fold higher (*P* ≤ 0.0165; one-way ANOVA; Fig. S1D), whereas doubling times did not differ significantly (Fig. S1E). Thus, Tween 80 was dispensable for growth but essential for optical clarity.

In IMDM, cell densities declined under the conditions tested (Fig. 2A). At 48 hours, the cell density in EBM supported a 325-fold higher density than IMDM (*P* = 0.003; two-way ANOVA; Table S2). Sodium thioglycolate increased IMDM cell density 1.8-fold at 24 hours (*P* = 0.009; Fig. S1F), but did not prevent subsequent decline. Published IMDM cultivation often begins near 10 cells/mL, which is approximately 100-fold above our starting density^24,34^. Attempts to recover *Blastocystis* using published soft-agar^21,22^ and solid-agar^32^ were also unsuccessful under our test conditions.

A common observation in published *Blastocystis* growth curves is that the initial seeding density strongly impacts growth and carrying capacity^24–26^. We therefore tested growth in EBM medium across a range of seeding densities. ST1 cultures seeded from 10^5^-10^3^ cells/mL exhibited exponential growth and reached densities exceeding 7×10^6^ cells/mL (Fig. 2B). Thus, EBM medium supported exponential growth across the densities tested, including inocula below those used in published liquid-culture protocols^35^.

Finally, we characterized growth of the four axenic and clonal isolates representing ST1–ST4. All four isolates grew in EBM medium, although their growth dynamics differed (Fig. 2C,D; Fig. S2A–C). At an initial density of 10^5^ cells/mL, three isolates had modeled carrying capacities exceeding 6×10^6^ cells/mL, whereas the ST3 isolate reached 1.4×10^6^ cells/mL. Estimated doubling times ranged from 6.2 hours for ST4 to 12.9 hours for ST3, and the average modeled lag time across isolates was 2.3 hours. The isolates also differed in cell size (Fig. 2C; Fig. S2D; Table S2). The average cell area of the ST3 isolate was 169 μm², compared with 106, 109, and 32 μm² for ST1, ST2, and ST4, respectively.

### Clonal Isolation Resolves Within-Subtype Strain Diversity in a Single-Host Culture

We sequenced a xenic ST1 culture that had previously been identified as ST1 by 18S rRNA gene PCR. PacBio long-read sequencing produced a *Blastocystis* genome assembly of 21.9 Mb, substantially larger than previously published ST1 assemblies, which range from 14-18 Mb. BUSCO analysis identified 25 duplicated single-copy genes, a high amount relative to other *Blastocystis* genomes, including the ST1 reference isolate Nand II (ATCC 50177; Fig. 3A). The assembly also contained multiple full-length 18S rRNA gene sequences. The lowest pairwise nucleotide identity among these sequences was 98.5%, within the range of variation reported for sequences assigned to the same ST^10^. The combination of a large assembly, excess duplicated single-copy genes, and divergent 18S rRNA gene sequences suggested together suggested unresolved closely related strains. To test this interpretation, we performed clonal isolation on the original culture and sequenced the resulting isolate using Illumina short reads. Its genome was 15.4 Mb, with six duplicated BUSCO genes, values comparable to published ST1 genomes (Fig. 3A). Multiple full-length 18S rRNA copies remained, but shared 99.9% identity. The combined reduction in assembly size, duplicated genes, and 18S diversity was consistent with recovery of a single strain from a multi-strain starting culture.

**Figure 3:**
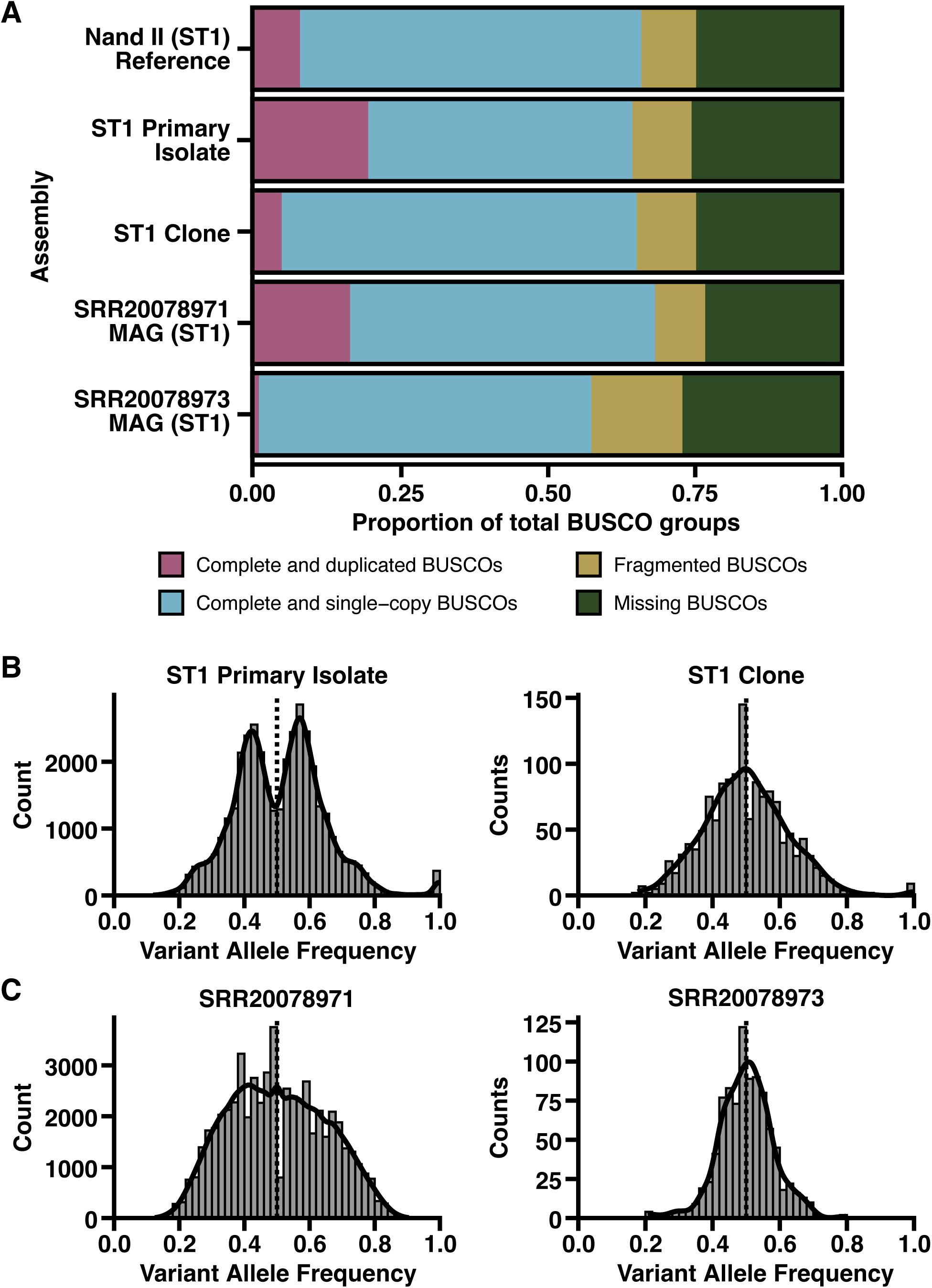
Clones are necessary for ST identification and assignment. **(A)** Proportions of BUSCO groups are shown for 5 *Blastocystis* ST1 assemblies generated from samples of a single strain and mixtures. **(B)** Allele frequency distributions of heterozygous variants of *Blastocystis* ST1 primary isolate and clone. Black curves represent the density curve of each distribution. Dashed lines represent allele frequencies of 0.5. **(C)** Allele frequency distributions of heterozygous variants of *Blastocystis* ST1 MAG. Black curves represent the density curve of each distribution. Dashed lines represent allele frequencies of 0.5.

To assess the mixture independently of assembly size, we mapped reads from both cultures to the ST1 Nand II genome and compared biallelic variant allele frequencies. In the derived isolate, heterozygous variants formed one peak near 0.5, as expected for a predominantly diploid, genetically homogeneous sample^11^(Fig. 3B). The original culture instead showed peaks near 0.42 and 0.58, with additional variants in the distribution tails. Together with the assembly and 18S results, this pattern supported the presence of at least two closely related ST1 strains in the original single-host culture.

### Signatures of Strain Mixtures in Metagenomic Sequencing Databases

To assess whether similar strain mixtures could be detected in previously sequenced samples, we screened public sequencing data for *Blastocystis* ST1. Using EukDetect2^8^, we screened over 36,000 human gut metagenomic samples and identified 91 samples with at least 1× coverage of the ST1 Nand II reference genome and no detected additional *Blastocystis* STs (Table S4)^13,36–40^. Only five of these samples had coverage greater than 10×, the minimum threshold we used for variant inference with DeepVariant^41^, limiting the number of metagenomic samples suitable for evaluating strain composition. We also examined four samples of genomic or metagenomic sequencing of cultures of *Blastocystis* ST1 taken from individual human donors, including three xenic cultures and one axenic culture^42,43^. All four had greater than 10x coverage depth of the Nand II reference genome.

With these samples, we inferred variants from read alignments and examined the resulting biallelic variant allele frequency distributions (Fig. S3). These distributions varied across samples, ranging from a single prominent peak near 0.5, consistent with a relatively homogeneous strain composition, to broader or multiple-peaked distributions consistent with strain mixtures.

Among deeply sequenced samples, two cultures from the same study^42^ clearly illustrated this contrast. Sample SRR20078973 showed a single prominent allele-frequency peak near 0.5, whereas SRR20078971 showed a broader distribution with multiple peaks (Fig. 3C). Genome assemblies from these samples showed a corresponding difference in duplicated BUSCO single-copy genes. The assembly from SRR20078973 contained one duplicated gene, whereas the assembly from SRR20078971 contained 21 (Fig. 3A). The concordance between allele-frequency distributions and BUSCO duplication in these independent cultures was consistent with a relatively homogeneous strain composition in SRR20078973 and a mixture of strains in SRR20078971.

Together, the analyses of our ST1 culture and previously sequenced cultures show that cultures identified as a single *Blastocystis* ST can contain genomic signatures consistent with multiple closely related strains.

## Discussion

By establishing a liquid, optically clear culture medium that supports exponential growth from low inoculum densities, EBM medium addresses a central limitation in *Blastocystis* experimental biology. EBM achieved higher modeled carrying capacities than biphasic egg medium and enabled direct screening of 96-well plates, time-lapse imaging, and modular axenization and clonal isolation. We recovered axenic and clonal isolates representing ST1–ST4, which are the subtypes most common in humans. Together, the medium and workflow make it feasible to expand collections of tractable isolates and compare their biology under controlled conditions. By supporting exponential growth at substantially lower inoculum densities, EBM may facilitate recovery of *Blastocystis* directly from primary samples, where protist abundance can vary^44^.

This work also establishes the importance of clonal isolation procedures in *Blastocystis* research. Here, a ST1 culture contained genome signatures of multiple closely related strains, including assembly size, single copy gene duplication, 18S sequence diversity, and allele-frequency structure. After clonal isolation, all signatures shifted toward the patterns expected for a single strain. An independent published ST1 culture from a human donor showed a similar combination of excess duplicated genes and complex allele frequencies.

These observations support within-ST strain coexistence in single-host-derived cultures, rather than merely co-occurrence of divergent subtypes. However, how frequently *Blastocystis* strains coexist within hosts remains unresolved. Only five of 91 qualifying human metagenomes provided sufficient coverage for our variant analysis, precluding a reliable prevalence estimate.

Importantly, single-subtype identification cannot be used as evidence of a single-strain culture. Here, conventional 18S genotyping did not reveal that the original ST1 culture contained a strain mixture. The most divergent full-length 18S sequences differed by approximately 1.5%, but commonly amplified diagnostic fragments span only 300–600 bp^45,46^ and may lack strain-discriminating sites. Sequencing the full-length 18S rRNA gene provides additional information but may not fully resolve strains as the extent of within-isolate 18S rRNA gene variation remains insufficiently characterized. Here, the clonal isolate contained full-length 18S rRNA sequences that were 99.9% identical. Genome-wide evidence and clonal isolation therefore provide important complementary means of assessing strain composition.

Resolving strain-level diversity is an import step for understanding *Blastocystis* biology. Closely related strains may differ in metabolism, interactions with other members of the gut microbiome, or effects on the host. Without clonal isolates, these properties cannot be attributed to a specific lineage, complicating comparisons across studies and cultures. EBM medium and our isolation pipeline provide a means to generate genetically defined cultures for testing these differences and to facilitate future genetic manipulation. Although genome-editing tools have not yet been established in *Blastocystis*, proof-of-concept studies have demonstrated plasmid delivery and expression^18,47^. Clonal isolation will be critical for recovering and characterizing genetically modified lines. More broadly, establishing clonality as part of isolate characterization may help ensure that phenotypes attributed to a particular *Blastocystis* ST or strain are interpreted against a defined genetic background.

EBM medium supports robust *Blastocystis* growth under conditions suitable for axenic and clonal isolation, providing a practical platform for experimental studies. Our findings also show that cultures identified as a single ST can contain genomic signatures consistent with multiple closely related strains. By facilitating the recovery of defined isolates, this approach enables more rigorous investigation of *Blastocystis* strain diversity, its ecological significance, and its potential contributions to host–microbiome interactions.

## Materials and Methods

### Sample collection

Fecal samples were obtained from nonhuman primates at Zoo Atlanta and from human participants (Table 1). Animal samples were collected under a protocol reviewed and approved by Zoo Atlanta’s Scientific Research Committee (SRC # 2024-16). Human samples were obtained from participants enrolled under a protocol approved by the Georgia Institute of Technology Office of Research Integrity and Assurance Institutional Review Board (IRB2025-772). Written informed consent was obtained from all participants prior to sample collection.

### Culture Conditions

*Blastocystis* cultivation and experimentation was conducted in an anaerobic chamber (Biolog Cat. No. AS-500) with an internal incubator at 36°C, filled with an anaerobic gas mixture (5% CO_2_; 5% H_2_; N_2_ balance). All media and plastic culture materials were pre-reduced inside the chamber for 24 hours prior to use to allow.

EBM medium contained Stone’s modification of Locke’s solution (ATCC medium 1671; Locke’s), chicken egg yolk, HIHS, sodium thioglycolate, and Tween 80 for optical clarity. Locke’s was autoclaved and sterile-filtered before addition of 4% (w/v) yolk and, optionally, 1% (w/v) Tween 80. After stirring (500 rpm, 60 min, room temperature), the mixture was filtered sequentially through a commercial coffee filter, Whatman grades 3 and 5 filter paper, and a 0.22-µm filter, then heated at 50°C for 60 min. The filtered mixture (75%, v/v) was combined with HIHS (25%, v/v) and sodium thioglycolate (0.5 mg/mL final) and re-sterile-filtered. For medium without Tween 80, components were filtered sequentially to minimize lipid precipitation.

Biphasic egg medium was prepared following ATCC medium 1671 with modifications: slants used unfertilized chicken eggs, autoclaved Stone’s modified Locke’s solution was sterile filtered, and 0.5 mg/mL sodium thioglycolate was added to the liquid phase.

IMDM (Thermo Fisher Cat. No. 12200036) was prepared per manufacturer instructions, supplemented with 10% v/v heat-inactivated horse serum (Thermo Fisher Cat. No. 26050088) with and without 0.5 mg/mL sodium thioglycolate.

### Blastocystis Isolation

Fecal samples were collected within 24 hrs of production. 5 mL of fecal material was homogenized in 40 mL of PBS (with 0.5 mg/mL sodium thioglycolate and pre-reduced), filtered through a 40 µm strainer, and washed by centrifugation at 700 × g for 5 min. The pellet was resuspended in PBS and the process repeated twice more. The final pellet was resuspended in 3 mL of PBS. A 1 mL aliquot was reserved as the unseparated fraction, while the remaining 2 mL was mixed with 2 mL of 80% Percoll and layered onto a 100-80-40-20% Percoll gradient. Following centrifugation at 1,000 × g, for 10 min, 200-500 µL from the gradient interfaces and unseparated fraction were inoculated into pre-reduced 24-well plates containing 1 mL EBM medium supplemented with antibiotic cocktail (100 μg/mL carbenicillin [Biomatik Cat. No. A2311]; 100 μg/mL streptomycin [Thermo Fisher Cat. No. 61224-0500]; 100 μg/mL vancomycin [Thermo Fisher Cat. No. J62790-06]; 50 μg/mL kanamycin [Thermo Fisher Cat. No. BP906-5], and 25 μg/mL amphotericin B (Sigma-Aldrich Cat. No. A2942).

To assess *Blastocystis* presence, DNA was extracted using the Qiagen PowerFecal Kit (Qiagen Cat. No. 51804) following manufacturer’s instructions. 18S PCRs were performed using primers BL18SPPF/BL18SR2PPR^48^ and Rdr5/BhRDr^45^ in GoTaq MasterMix (Table. S5). PCR products were sequenced with Sanger sequencing via Eurofins and ST designation was determined by using BLASTn on the NCBI BLAST.

### Blastocystis Cultivation

Axenic and xenic *Blastocystis* cultures were grown at 36°C inside an anaerobic chamber (see anaerobic conditions section above) in 25 cm^2^, vented, tissue culture treated flasks (Thermo Fisher Cat. No. FB012935) that were positioned upright. Each culture was maintained in 20 mL of EBM medium, and passaged by making 1:20-1:200 dilutions into fresh media every 3-4 days depending on ST specific growth rates. Xenic cultures were passaged with an antibiotic cocktail as described above. Cells were resuspended by gentle swirling prior to use.

### Growth Assays

Triplicate cultures were seeded at 10□ cells/mL without antibiotics in flasks containing 20 mL EBM medium or IMDM, or tubes containing biphasic egg medium (6 mL liquid, 4.5 mL slant). At each timepoint, cultures were resuspended and 100 µL aliquots removed and counted. Plate-based growth curves used independent triplicate cultures seeded at 10³–10□ cells/mL in 1 mL EBM per well. Growth curves were fitted using a logistic model incorporating a Heaviside step function for lag time. Statistical analyses were performed in Prism v10.6.1. Cell densities were log10-transformed, with zero values assigned half the detection limit value. Growth curves were compared using two-way mixed-effects ANOVA with Tukey-adjusted comparisons within timepoints. Growth parameters were analyzed without transformation using unequal-variance one-way ANOVA with Dunnett-adjusted comparisons.

Endpoint growth assays were seeded at 10□ cells/mL in 24-well plates and counted after 48 hrs at 36°C. Conditions were compared using unequal-variance one-way ANOVA with Dunnett-adjusted comparisons.

### Live Cell Microscopy and Analysis

Images were acquired using a Nikon Ts2 inverted microscope with a Kiralux 5.0 MP CMOS camera (Thorlabs Cat. No. CS505CU1). Cell areas were quantified in FIJI/ImageJ v1.54p by thresholding the top 1% of pixel intensities and analyzing particles with areas of 50–8000 pixels and circularity of 0.2–1.0. Statistical comparisons were performed in Prism v10.6.1 using unequal-variance one-way ANOVA assuming log-normal distributions, with Games–Howell-adjusted comparisons.

### Clonal Isolation Protocol

*Blastocystis* cultures were grown antibiotic-free in EBM medium for 48 hrs before dilution to 25 cells/mL in 50 mL fresh EBM medium. 100 μL/well was added to 96-well plates, sealed in plastic bags to minimize evaporation, and incubated at 36°C for ≥5 days. Wells were screened microscopically with a 4X objective (see microscopy details), limiting aerobic atmospheric exposure to 15 min, and incubated an additional 48 hrs if no growth was detected. Wells containing only single visible foci were expanded in 1 mL EBM medium in 24-well plates for 2–5 days, then transferred to flasks containing 20 mL EBM medium for routine cultivation. Wells containing multiple colonies were excluded.

### Clonal Confirmation

Candidate clonal cultures were preliminarily evaluated for clonality by full-length 18S PCR and Nanopore sequencing (Plasmidsaurus). 50 µL culture aliquots were lysed in 200 µL DNAzol Direct (20 mM potassium hydroxide; 60% [w/v] PEG 200, pH 13.3–13.7) at 80°C for 10 min. PCR reactions with SSU-F1/SSU-R1 primers^49^ were conducted using GoTaq MasterMix (Table S5). Full-length sequences were assigned ST by BLAST and required ≥ 98% within-sample pairwise identity.

### Axenization Protocol

*Blastocystis* cultures were grown in EBM medium in flasks for 48 hrs with the antibiotic cocktail described above. Then 50 mLs of fresh EBM medium with 25 μg/mL levofloxacin (from 50 mg/mL stock dissolved in DMSO [Thermo Fisher Cat. No. J66943.03]) were seeded with 25 cells/mL and grown and screened as above. Candidate wells were pooled into 20 mL antibiotic-free EBM medium and grown for 5–7 days or until visible growth. Pooling improved survival following levofloxacin exposure but precluded clonal isolation.

Candidate axenic cultures were screened for bacterial growth by inoculating 5 µL onto Brucella blood agar (Biolog, AS-141) and Yeast Casitone Fatty Acids Agar with Carbohydrates (Biolog, AS-675), followed by anaerobic incubation at 36°C for 48 hrs. Cultures without visible bacterial growth were further assessed by 16S rRNA PCR. 50 µL culture aliquots were treated with 1 µL Turbo DNase (Thermo Fisher, AM2238) at 37°C for 1 hr to remove extracellular DNA, then lysed with 200 µL DNAzol Direct at 80°C for 10 min. PCR reactions used 27F/1492R primers^50^ in GoTaq MasterMix (Table S5). PCR products were visualized in 1% agarose gels.

### Blastocystis ST1 DNA extraction and sequencing

DNA from xenic, nonclonal *Blastocystis* ST1 and axenic, clonal ST1 cultures as previously described^51^ and assessed by Qubit fluorometer, Agilent Genomic DNA Tapestation, and Nanodrop. The original culture was sequenced at Maryland Genomics at the University of Maryland (Baltimore, Maryland, USA) on a PacBio SMRT cell run on a PacBio Revio, generating 750,660 reads with a median read N50 of 11,048 bp. The derived isolate was sequenced at SeqCenter (Pittsburgh, PA, USA) using Illumina 150 bp paired-end reads and sequenced on an Illumina NovaSeq X Plus.

Xenic, non-clonally isolated *Blastocystis* ST1 PacBio long reads were assembled using hifiasm-meta hamt version 0.3-r073^52^. Eukaryotic contigs were identified using MetaEuk taxtocontig version 1f7d7023bdac3d46964ca380bef1b2848685501e^53^ and a custom protein database combining the Swissprot reference database with protist proteomes available through UniProt. Axenic, clonally isolated *Blastocystis* ST1 Illumina paired-end short reads were assembled with SPAdes version 4.2.0^54^ using the --isolate flag. 18S rRNA gene regions were identified in each genome by running blastn version 2.16.0 on a *Blastocystis* ST1 18S rRNA full length gene (GenBank accession KY610205.1). Average nucleotide identity between full length 18S genes was calculated using the emboss version 6.0.0 needle algorithm. BUSCO^55^ version 6.0.0 was used for single copy gene analysis, run with the eukaryota_odb12 lineage database.

### Public metagenome analysis and variant calling

Over 36,000 public metagenomes were analyzed for the presence of *Blastocystis* using EukDetect2^8^.

Samples containing only *Blastocystis* ST1 and no other STs were retained. Those with high estimated absolute abundances (RPKSB) were aligned to the *Blastocystis* ST1 Nand II reference (GCA_039096935.1) using minimap2 2.30^56^ and coverage depth was estimated using samtools^57^ version 1.23. Alignments within 2 kb of contig ends were excluded, and mapping quality ≥30 was required. Samples with greater than 1x average genome coverage (Table S4) were variant-called with DeepVariant^58^ in the appropriate whole-genome or PacBio mode, then filtered with BCFtools 1.23.1^59^ for variant and genotype quality ≥20. Phased and unphased heterozygous biallelic sites frequencies were plotted in R.

## Data, Metadata, and Code Availability

Sequencing data from the xenic multi-strain ST1 isolate and the axenic single strain ST1 isolate are available on the NCBI Short Read Archive under BioProject PRJNA1537639. *Blastocystis* contigs assembled from both sequencing datasets are available on Zenodo at https://doi.org/10.5281/zenodo.23265426. Code used for the variant calling and variant analysis are available on GitHub at https://github.com/jshih08/blasto_clonal_analyses and archived at Zenodo at https://doi.org/10.5281/zenodo.23267582. All public data used in the study is reported in Table S4.

## Supporting information

Supplemental Movie 1: Timelapse of Blastocystis growth in monophasic media

Supplemental Table 1: Axenic strains at ATCC

Supplemental Table 2: Raw in vitro data

Supplemental Table 3: ST metadata

Supplemental Table 4: Coverage of Blastocystis ST1 samples

Supplemental Table 5: Primers and PCR parameters

## Acknowledgements

A.L. and F.L. conceptualized the project. A.L., F.L., J.S. and L.K. designed experiments. F.L., J.S., L.K., A.L., S.P., L.D., and G.N. conducted experiments. F.L., A.L., J.S., and L.K. wrote and revised the manuscript.

This work was funded by NIAID K22 award K22AI173181, NIGMS R35GM166073, and Georgia Institute of Technology to A.L., by NIAID postdoctoral fellowship F32AI200531 to F.L., and by T32GM142616 to J.S. The funders had no role in study design, data collection and interpretation, or the decision to submit the work for publication.

We thank Zoo Atlanta and Joseph Mendelson for help with sample collection from non-human primate samples. We also thank Lauren Speare and Nathan Mcdonald for helpful feedback.

## Figures

**Supplemental Figure 1:**
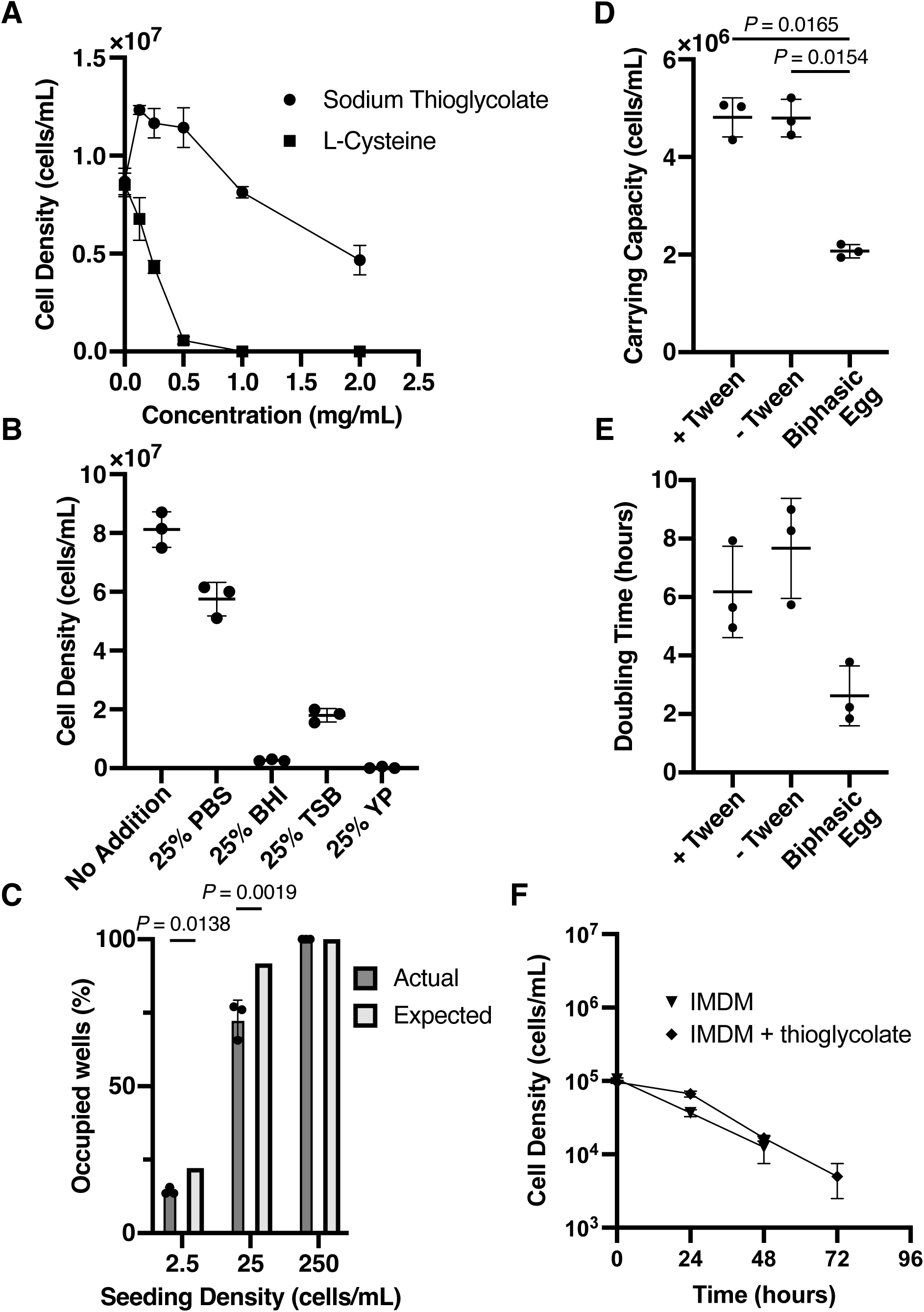
Media formulation tests and growth curve dynamics. **(A)** Sodium thioglycolate and L-cysteine reducing agents impact on *Blastocystis* ST1 cultures. Cell density of *Blastocystis* ST1 cultures after 48 hours of growth seeded at 10^5^ cells/mL in EBM media with no reducing agent added or varying concentrations of sodium thioglycolate or L-cysteine. The monophasic media formulation includes 0.5 mg/mL of sodium thioglycolate. **(B)** *Blastocystis* does not tolerate common anaerobic bacterial growth media. Cell density of *Blastocystis* ST1 culture after 48 hours in EBM medium with or without 25% supplemented media. The control condition is PBS, phosphate buffered saline: BHI, brain heart infusion media: TSB, tryptic soy broth media: YP, yeast peptone media. All pairwise comparisons have *P*-values < 0.05, determined by one-way ANOVA. **(C)** *Blastocystis* survival decreases at low cell densities. The percentage of *Blastocystis* ST1 occupied wells compared to the expected occupation frequency as determined by poisson distribution. *P*-values were calculated by chi-squared test. **(D)** Carrying capacity of *Blastocystis* in EBM medium. Parameters extracted from Fig. 2A, excluding IMDM data since the same model could not be fit to declining population densities. *P*-values were calculated by one-way ANOVA and only values below 0.05 are displayed. **(E)** Doubling time of *Blastocystis* in EBM medium. Parameters extracted from Fig. 2A, excluding IMDM data since the same model could not be fit to declining population densities. *P*-values were calculated by one-way ANOVA and no conditions were significantly different. **(F)** *Blastocystis* growth in IMDM with and without sodium thioglycolate. Data for IMDM (without sodium thioglycolate) is from Fig. 2A, paired with growth data from IMDM with sodium thioglycolate. At 24 hours IMDM with sodium thioglycolate has a 1.8-fold higher cell density (*P* = 0.009, two-way ANOVA). No other time points are significantly different.

**Supplemental Figure 2:**
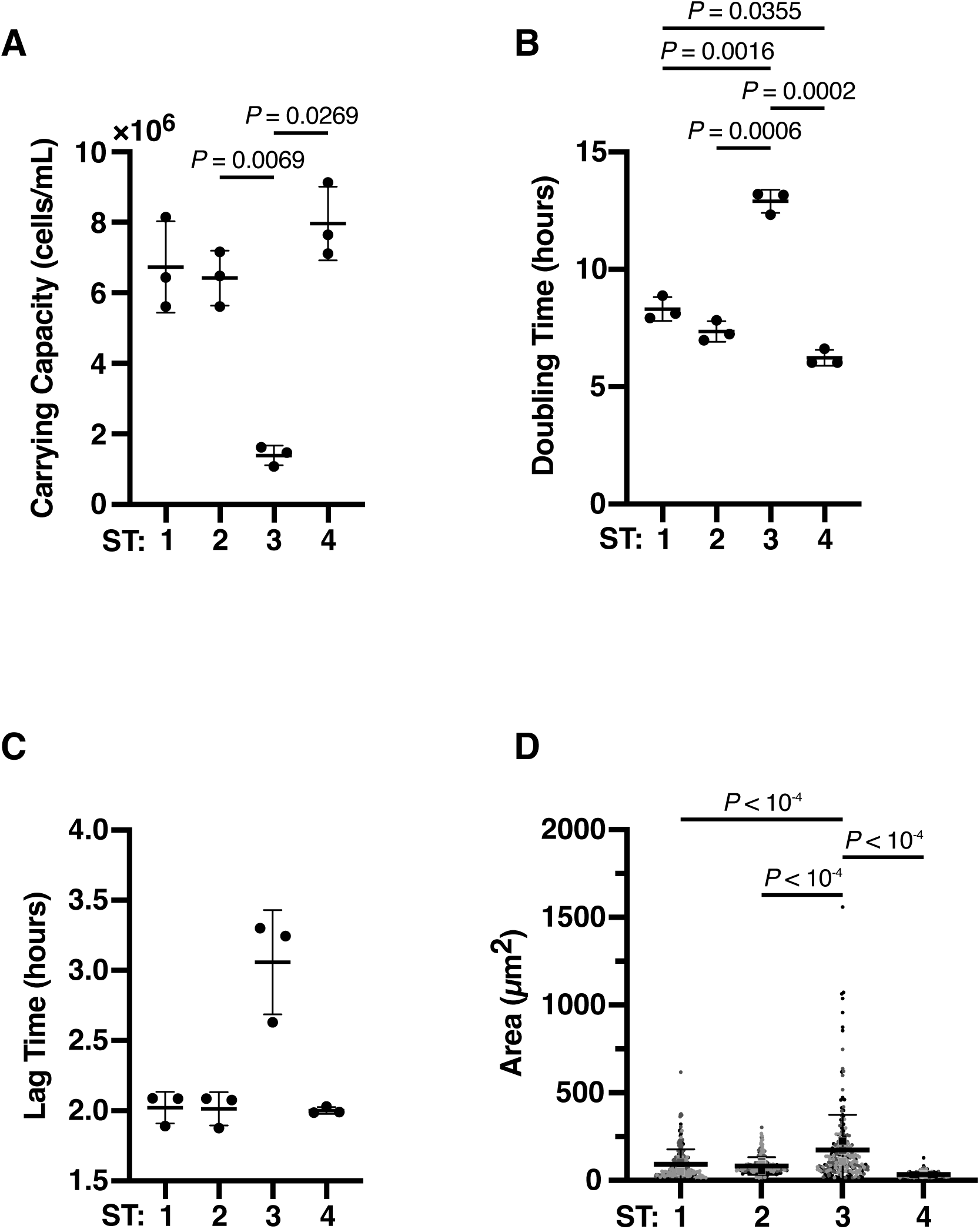
Differences in *Blastocystis* STs growth and size.(A) The carrying capacity of STs 1-4, determined by data from Fig. 2D. *P*-values were calculated by one-way ANOVA **(B)** The doubling time of STs 1-4. *P*-values were calculated by one-way ANOVA **(C)** The lag time of STs 1-4. No *P*-values were significant. **(D)** The cell size of STs 1-4. Cell sizes correspond to the area occupied by each cell in 2D images. Displayed *P*-values are only for comparisons to ST3 to demonstrate larger cell size. *P*-values were calculated by one-way ANOVA

**Supplemental Figure 3:**
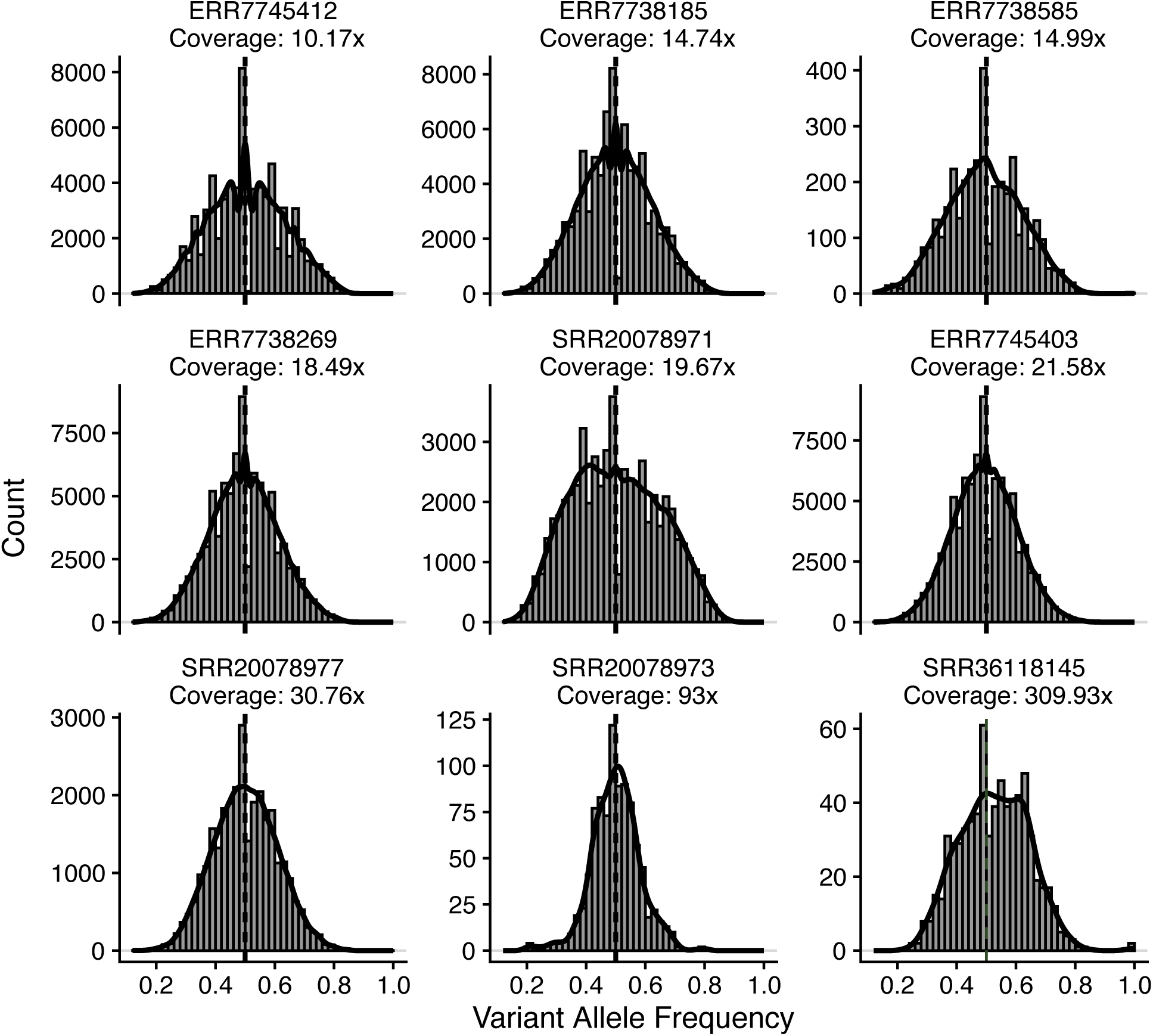
**Allele frequency distribution of public *Blastocystis* ST1 samples at high coverage. Black lines represent the density curve of each distribution. Dashed lines represent allele frequencies of 0.5.**

**Supplemental Movie 1:** Timelapse of *Blastocystis* growth in monophasic media

**Supplemental Table 1:** Axenic strains at ATCC

**Supplemental Table 2:** Raw *in vitro* data

**Supplemental Table 3:** ST metadata

**Supplemental Table 4:** Coverage of *Blastocystis* ST1 samples

**Supplemental Table 5:** Primers and PCR parameters

