## Supplementary figures and images for "A novel clonal and axenic isolation pipeline for the human gut protist *Blastocystis* reveals within-subtype strain coexistence"

### Supplemental Movie 1: Timelapse of Blastocystis growth in monophasic media

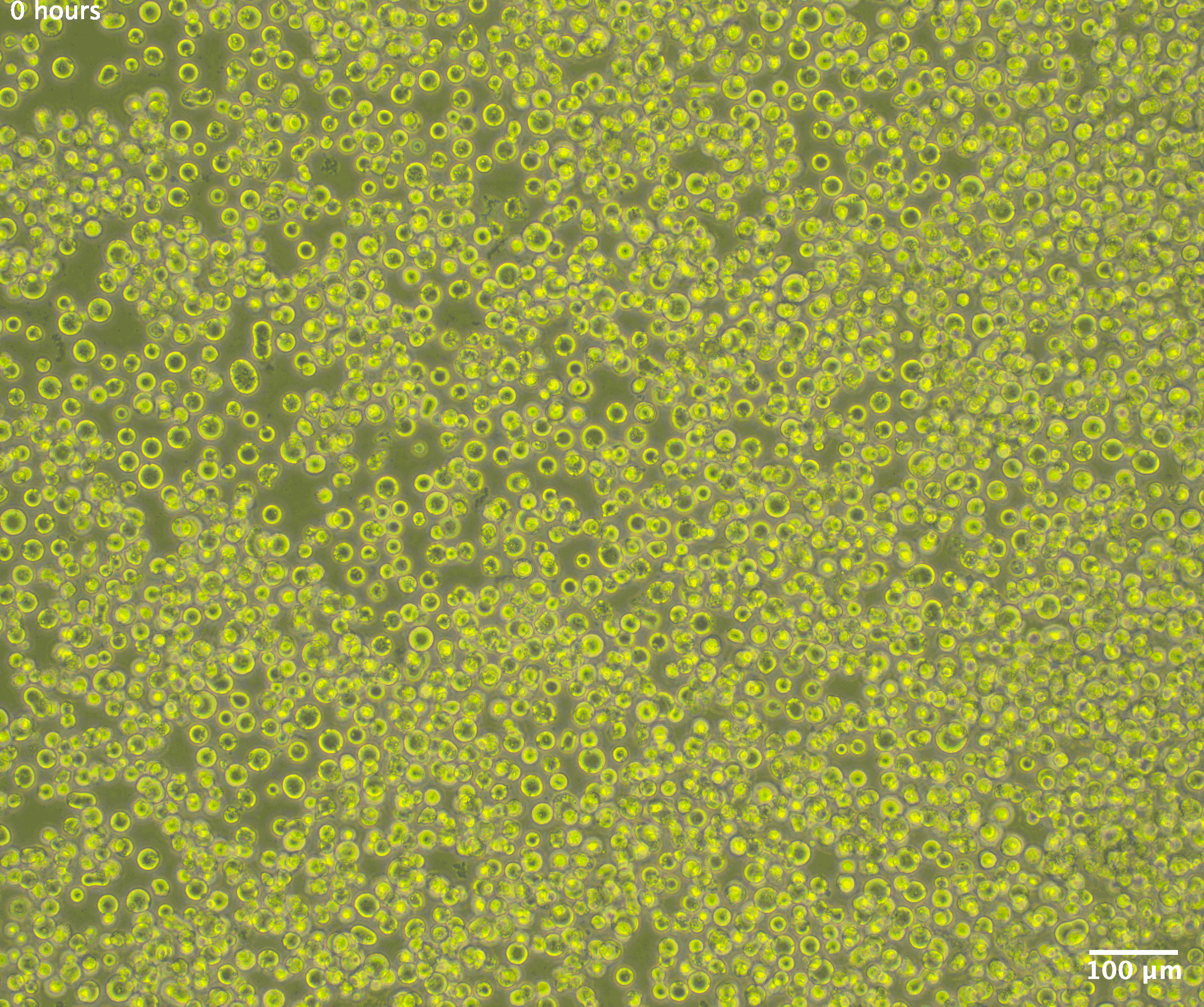
